# An efficient method to generate fluorescent amyloid fibrils

**DOI:** 10.1101/2022.12.28.522083

**Authors:** Kailash Prasad Prajapati, Masihuzzaman Ansari, Deepak Kumar Yadav, Bibin Gnanadhason Anand, Shikha Mittal, Karunakar Kar

**Author notes:** Authors have equal contribution (Kailash Prasad Prajapati, Masihuzzaman Ansari). Corresponding author, Karunakar Kar; School of Life Sciences, Jawaharlal Nehru University, New Delhi-110067, India.

## Abstract

Studies on fluorophore-tagged peptides help in elucidating the molecular mechanism of amyloidogenesis including their cellular internalization and crosstalk potential. Despite several advantages, unavoidable difficulties including expensive and tedious synthesis-protocols exist in fluorophore-based tools. Importantly, covalently-tagged fluorophores could introduce structural constraints which may influence the conformation of the monomeric and aggregated forms of protein. To resolve this problem, we describe a robust yet simple method to make fluorescent amyloid fibrils through non-covalent incorporation of fluorophores into amyloid fibrils. We used aggregation protocol in which a small amount of fluorophore is incorporated into the amyloids, and this protocol does not alter the aggregation kinetics and the characteristic β-sheet-conformers of the generated amyloid fibrils. We have successfully prepared fluorescent amyloid fibrils of Insulin, Lysozyme and Aβ_1-42_, and the noncovalently incorporated fluorophores remained intact in the amyloid fibrils without leaching, even after serial-dilutions and prolonged-storage. Further, this method enables successful monitoring of cellular-internalization of the fluorescent amyloids into SH-SY5Y and A549 cells, and it also detects FRET-signals during interfibrillar interactions. The findings establish a simple and affordable protocol to prepare fluorescent amyloid structures, which may significantly help amyloid researchers working on both *in vitro* and animal model systems.

## INTRODUCTION

Formation of amyloid fibrils due to aggregation of proteins and peptides has certainly gained huge attention, because the amyloid formation is directly linked to several complications including devastating neurodegenerative diseases.[1–3] Hence, it becomes important to understand various biophysical properties of the amyloids and their toxic effects on vital cellular activities.[4–6] Use of fluorophore-tagged peptides and proteins has been facilitating the mechanistic understanding of the internalization and the localisation of amyloid aggregates and their intervention with crucial cellular metabolism as well [4, 5, 7, 8]. Usually, a fluorescent amyloidogenic peptide is synthesized by tagging the desired fluorophore at an appropriate position in the peptide backbone or side chains through a convenient covalent bond [6, 7, 9, 10]. For example, thiol group mediated cross-linking reactions, preferably using cysteine residues in the peptide sequence [11]. In some cases, the direct covalent attachment of suitable fluorophores to the peptides is achieved during chemical synthesis of peptides[12, 13]. Though use of fluorophore-tagged peptides has become a vital tool for amyloid research, this protocol is not free from complications including its high cost and the difficulties in both the synthesis and the yield of the desired peptides. More importantly, the presence of the fluorophore molecule, as an inbuilt moiety in the polypeptide chain, may influence the inherent properties of the proteins including their actual conformation, their aggregation propensity, and the architecture of their amyloid assembly[8–10]. Due to the existence of the above-mentioned complications with the covalent tagging of fluorophores with peptides, it is very important for development of an affordable and simpler method for making fluorescent amyloid structures.

Recent research has revealed that aromatic metabolites are inherently aggregation-prone and their self-assembly can result in the formation of cytotoxic amyloid-like higher order structures[14–19]. Co-assembly of aromatic compounds have also been recently reported and such process was found to be accelerated in the presence of pre-formed amyloid structures[16, 17, 20]. For example, amyloid fibrils of Aβ_1-40_ is known to induce co-aggregation of the aromatic amino acids[20]. On the other side, amyloid-like structures of the metabolites are known to trigger aggregation of different globular proteins through effective amyloid cross-seeding[14–17, 21]. All these important revelations on the occurrence of the amyloid cross-seeding and co-aggregation between aromatic metabolites and proteins inspired us to hypothesize that, the addition of the aromatic fluorophores to a protein solution undergoing aggregation may possibly result in their co-assembly event, and if so, the end products of such process could turn out to be fluorophore-incorporated amyloid aggregates. To test this hypothesis, we conducted *in vitro* amyloid formation reactions of selected globular proteins as well as Aβ peptides, considering them as convenient amyloid systems. Rhodamine B was selected as the fluorophore for this study. Rhodamine-B (Rho-B) has intrinsic fluorescence property that has a characteristic λ_max_ of 575 nm when excited at 555 nm [22, 23]. The covalent attachment of Rhodamine-B to the proteins has been used to visualize the internalization of the proteins into the tissues and cells as well [24, 25]. Proteins with covalently tagged Rhodamine-B have also been used to study protein-protein interactions[24, 25].

Here, we report an innovative aggregation protocol for proteins which can yield the formation of fluorescent amyloid fibrils. Various biophysical and computational methods have been conducted to validate this simple method for preparation of the fluorescent amyloid fibrils through noncovalent incorporation of a small amount of the fluorophore component. This method is based on the fundamental principles of co-assembly during amyloid formation and only traces of the fluorophore molecules were added to the protein solution during the onset of the aggregation process. Hence, there is negligible perturbation to the structural and functional properties of the resultant β-sheet-rich amyloid fibrils, making them convenient amyloid systems for both cellular and *in vitro* experiments.

## EXPERIMENTAL SECTION

### Reagents

All Proteins and fluorophores were procured from Sigma-Aldrich and concentration was measured using Shimazdu uv-1900 spectrophotometer. Extinction coefficient values used as follows: 6080 M cm at 278 for insulin, 36 mM.cm at 280 nm for lysozyme, 43824 M cm at 280 nm for BSA, 106,000 M CM at 555 nm for Rhodamine-B, 76,900 M cm at 490 nm for Fluorescein.

### Sample preparation

Stock solutions were prepared as follows: Stock solution of Rhodamine-B was prepared by dissolving in PBS buffer (pH-7.4) & Fluorescein was prepared by dissolving in 0.1 M tris-HCl (pH-8.0). Stock solution of proteins were prepared by dissolving into PBS buffer, pH-7.4.

### Aggregation studies of proteins in presence of Rhodamine-B and Fluorescein

Thioflavin T, a fluorescent dye that specifically binds to amyloids aggregates[26], was used for aggregation studies. Aggregation kinetic studies of protein samples in the presence and absence of fluorophores were performed in PBS buffer, pH 7.4 at 65 °C. ThT concentration was maintained at 30 μM, and the fluorescence intensity of ThT at different time points was recorded using a Shimazdu fluorescence spectrophotometer (RF-5300, Japan). The rise in ThT was detected at 490 nm by exciting the ThT molecule at 440 nm. Aggregation reaction of all protein samples were carried out at 100 μM for insulin, 50 μM for lysozyme, with Rhodamine-B and Fluorescein at molar ratio of 1:100, 1:20, and 1:10.

### Incorporation of fluorophores into Protein fibrils

Fluorophore absorbance spectra of the supernatant of aggregating protein solutions having fluorophore at different time points was recorded using Shimazdu uv-1900 spectrophotometer, after centrifugation at 15,000 rpm for 10 minutes. Next, washing of the pellet was performed to remove excess fluorophores in the soluble form. For this, all the solutions after 72 hours were washed three times using PBS buffer, each wash involving centrifugation at 15000 rpm for 10 min and resuspension of pellet into 1 ml PBS buffer. For stable incorporation of fluorophore into protein aggregates again after 2-day centrifugation of solution was done and absorbance spectra of supernatant was obtained, and pellet was resuspended into 1ml PBS buffer. Detailed protocol is given in supplementary information.

### Fluorescence microscopy

The fluorescence microscopy was conducted using a fluorescence microscope (Nikon). 20 μl diluted aggregate sample was smeared over a glass slide, dried, and then stained with ThT and imaged, FITC filter was used for ThT imaging and TRITC used for Rhodamine-B and Fluorescein imaging. The cellular imaging studies were also performed using the same Nikon 90i microscope.

### Cell Culture studies

The human neuroblastoma cell line (SH-SY5Y) and Lung cancer A549 cell lines were obtained from the American Type Culture Collection (ATCC, CRL-2266). The cells were grown in DMEM/ F12 (Gibco Life Technologies, USA) medium supplemented with 10% heat inactivated fetal bovine serum (FBS) (Gibco Life Technologies, USA) and 50 U/ml penicillin, 100 μg/ml streptomycin (HIMEDIA). Cells were maintained in an incubator at 37°C with 5% atmospheric CO_2_ and 95% humidity. The medium was changed after 48 h. Description of methods for MTT assay and live cell imaging studies (confocal and fluorescence microscopy) is given in supplementary information.

## 3. RESULTS AND DISCUSSION

### Preparation of the fluorescent insulin amyloids

The study begins with amyloid aggregation reactions of the insulin solution in physiological buffer conditions (PBS, at 65 °C) by using amyloid specific Thioflavin T assay [26]. The concentration of the insulin was chosen following the previously established studies[27, 28]. The rise in the ThT signal for [100 μM insulin + 5 μM Rho-B] sample was observed to follow the same pattern as observed for the control insulin sample at 100 μM in the absence of the fluorophore (Figure 1a 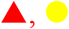). This result suggests that the presence of RhoB in the insulin solution (at 100:5 molar ratio of Insulin:RhoB) does not alter the aggregation kinetics of the insulin. ThT signal of the control solution containing only Rho-B at 5μM remained unchanged throughout the studied time frame which confirmed that the Rho-B alone (at the studied concentrations) does not aggregate, and its presence does not interfere with the ThT fluorescence (Figure 1b, 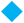). The important question was to check whether Rho-B molecules are incorporated into the resultant insulin aggregates at the end of the reaction. To verify this, we compared the absorbance spectra of the [ins + Rho-B] sample before aggregation at 0 h and after aggregation (supernatant of the final aggregate suspension obtained at 8 d). The obtained data as shown in **Figure 1b** clearly revealed substantial reduction in the Rho-B absorption peak, suggesting the incorporation of Rho-B into insulin fibrils. Correlation between increased ThT-signal of the (Ins+Rho-B) sample and the decreased Rho-B concentration in the supernatant of the centrifuged aggregated suspension at different time points is shown in **Figure 1c** which verify the direct incorporation of Rho-B into the insulin amyloid conformers during the progression of insulin fibril formation. After establishing the direct incorporation of RhoB into insulin amyloid fibrils, we tried different combination of the Insulin:RhoB molar ratio. For example, at 100:1 molar ratio of Insulin: RhoB, direct incorporation of the RhoB into the insulin aggregates was observed without affecting the aggregation kinetics of insulin (**Figure 1e,1f**). Similar results were obtained for the sample containing 100:10 molar ratio of insulin:RhoB (**Figure 1g,1h**).

**Figure 1.**
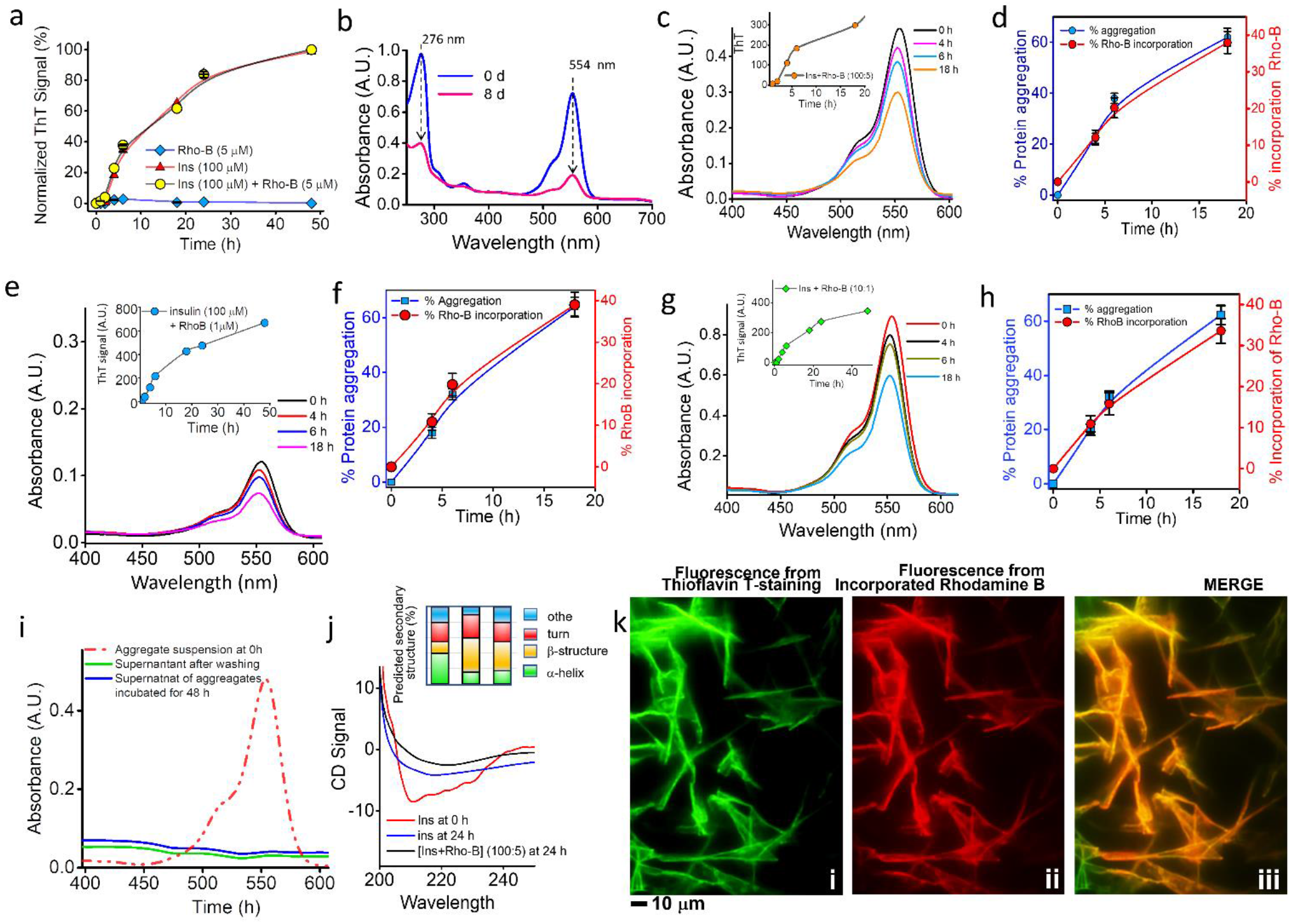
Noncovalent incorporation of Rhodamine B into insulin amyloid fibrils. ***a,*** ThT assays on temperature-induced amyloid formation of 100 μM insulin in PBS, pH 7.4: 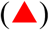 Ins only; 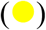 Ins + Rho-B (at 100:5 molar ratio); 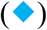 RhoB alone. ***b,*** Rho-B absorbance in supernatant of aggregated insulin solution at 0 h 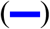 and after 8 d 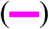. ***c,*** Decrease in the Rho-B absorbance in the supernatant of centrifuged [100 μM Ins + 5 μM Rho-B] sample during aggregation at different time points. Inset is the rise in the ThT signal. ***d,*** Correlation between extent of aggregation and the decrease in Rho-B concentration in the supernatant, confirming incorporation of Rho-B into insulin aggregates. ***e,*** Decrease in the Rho-B absorbance in the supernatant of [insulin + Rho-B, at 100:1 molar ratio]. Inset shows the respective rise in the ThT signal. ***f,*** Correlation between % aggregation of Ins the decrease in the Rho-B absorbance for [ins + Rho-B, at 10:1] sample. ***g,*** Rho-B absorbance in the supernatant of [100 μM ins + 10 μM Rho-B] sample during aggregation. Inset shows rise in the ThT signal. ***h,*** Correlation between % ins aggregation and the decrease in the Rho-B absorbance for [100 μM ins + 10 μM Rho-B] sample. ***i,*** Absorbance of Rho-B in (Ins+Rho-B) sample at different conditions: 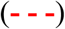 supernatant at 0 h before aggregation; 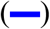 supernatant of the centrifuged resuspended [ins+Rho-B] aggregates; 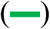 supernatant of the centrifuged [ins+Rho-B] aggregate suspension after 2 d storage. ***j,*** CD data show the β-sheet rich structures for ins amyloid aggregates after Rho-B incorporation: 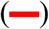 ins at 0 h; (**—**) ins at 48 h; 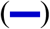 [ins +Rho-B] aggregates at 48 h. Inset shows the % predicted secondary structures. ***k,*** Fluorescence microscopy of the Rho-B incorporated Insulin fibrils: (i) ThT-stained (external dye); (ii) Intrinsic fluorescence of Rho-B without any external dye; (iii) Merged view revealing the internalization of Rho-B into amyloid fibrils.

To check whether the Rhodamine B is leaching out of the fibrils due to dilution or prolonged storage, we examined the absorption spectra of the supernatant of the centrifuged RhoB-incorporated insulin-fibril suspension which was stored for 4 weeks. The data shown in **Figure 1i** confirmed the absence of RhoB peak in the supernatant. Similarly, after serial dilutions, we did not see any leaching of the RhoB dye from the insulin fibrils. These results confirmed that RhoB can be incorporated into the insulin amyloid fibrils noncovalently and these incorporated RhoB molecules remain intact within the fibrillar architecture without leaching out. Non-covalent incorporation of RhoB into insulin fibrils is further confirmed by our disassembly experiments in which the release of free RhoB from insulin fibrils were detected after papain-mediated disaggregation reaction[29] (**Figure 2**). This result validates two important usefulness of this method: 1) the incorporation of RhoB into amyloid fibril; 2) Monitoring the released RhoB can be used as a tool for studying the amyloid disassembly process.

**Figure 2.**
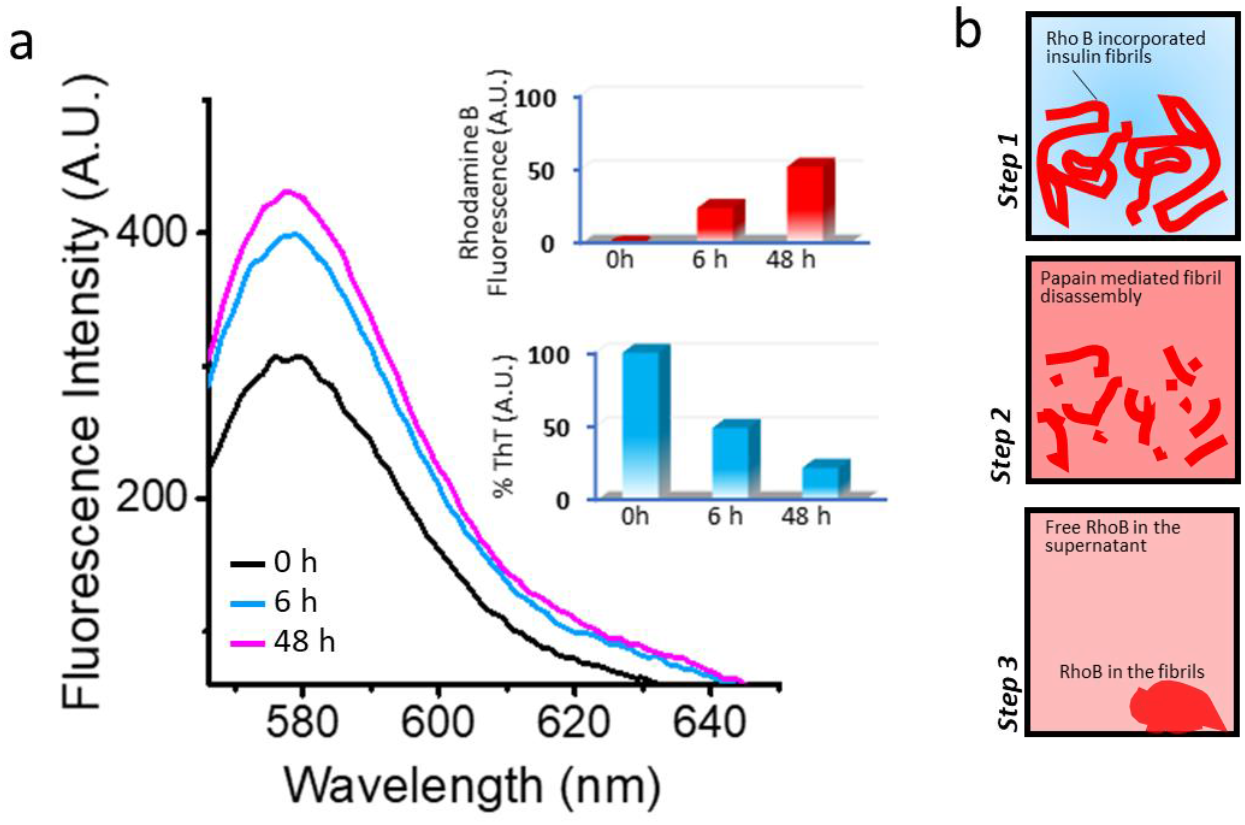
Study of amyloid disassembly via fluorescence of released Rhodamine B. Fluorescence emission of supernatant of centrifuged sample of RhoB-incorporated insulin fibril suspension, revealing release of free Rhodamine B molecules from the fibrillar structures, a different time point (0h, 6h and 48 h). Inset upper panel shows gradual increase of RhoB emission (at 575 nm) in the supernatant. Inset lower panel shows gradual decrease of the ThT signal of the aggregate suspension, confirming the disaggregation of amyloid structures. B. Schematic presentation of the protocol for disassembly assay on RhoB-incorporated amyloid fibrils.

Next step was to visualise RhoB-incorporated insulin fibrils using fluorescence microscopy, and our images revealed the presence of RhoB-specific red fluorescent fibrils (Figure 1 k panel ii). The same sample displayed green-fluorescent fibrils after ThT staining (**Figure 1 k, panel i**) and both fluorescence signals appeared to overlap with each other in the merged view (**Figure 1 k, panel iii**). To examine the secondary structures of the insulin fibrils incorporated with RhoB, we performed CD experiments, and the obtained data clearly suggest the prevalence of β-sheet rich structures in both control insulin fibrils and RhoB incorporated insulin fibrils (**Figure 1j, inset**). Based on these results out protocol enables successful incorporation of RhoB into the fibrillar architecture of insulin without altering characteristic of the β-sheet rich structure of resultant amyloid fibrils as well as aggregation reaction kinetics.

We further extended this study to generalise on non-covalent incorporation of RhoB into amyloid fibrils. We applied this protocol to make fluorescent amyloid of lysozyme and Aβ_1-42_ peptide **Figure 3**. As described method in above for making RhoB incorporated insulin fibers, we successfully prepared RhoB incorporated Lysozyme amyloid fibers **Figure 3 a**. The embed rhodamine b in lysozyme fibers remained intact without leaching from the fibril as evident from **(Figure 3 b)**. The secondary structure obtained from CD experiment of lysozyme amyloid fibrils incorporated with RhoB have similar beta structure as observed for control lysozyme fibrils **(Figure 3 c, inset).** Next step was to visualise RhoB-incorporated lysozyme fibrils using fluorescence microscopy, and our images revealed the presence of RhoB-specific red fluorescent fibrils **(Figure 3 d panel ii)**. The same sample displayed green fluorescent fibrils after ThT staining (**Figure 3 d, panel i**) and both fluorescence signals appeared to overlap with each other in the merged view (**Figure 3 d, panel iii**). Next, we have extended this method for non-covalent incorporation of RhoB into Alzheimer’s disease linked amyloid fibril formation of Aβ_1-42_ peptides **(Figure 3 e)**.

**Figure 3.**
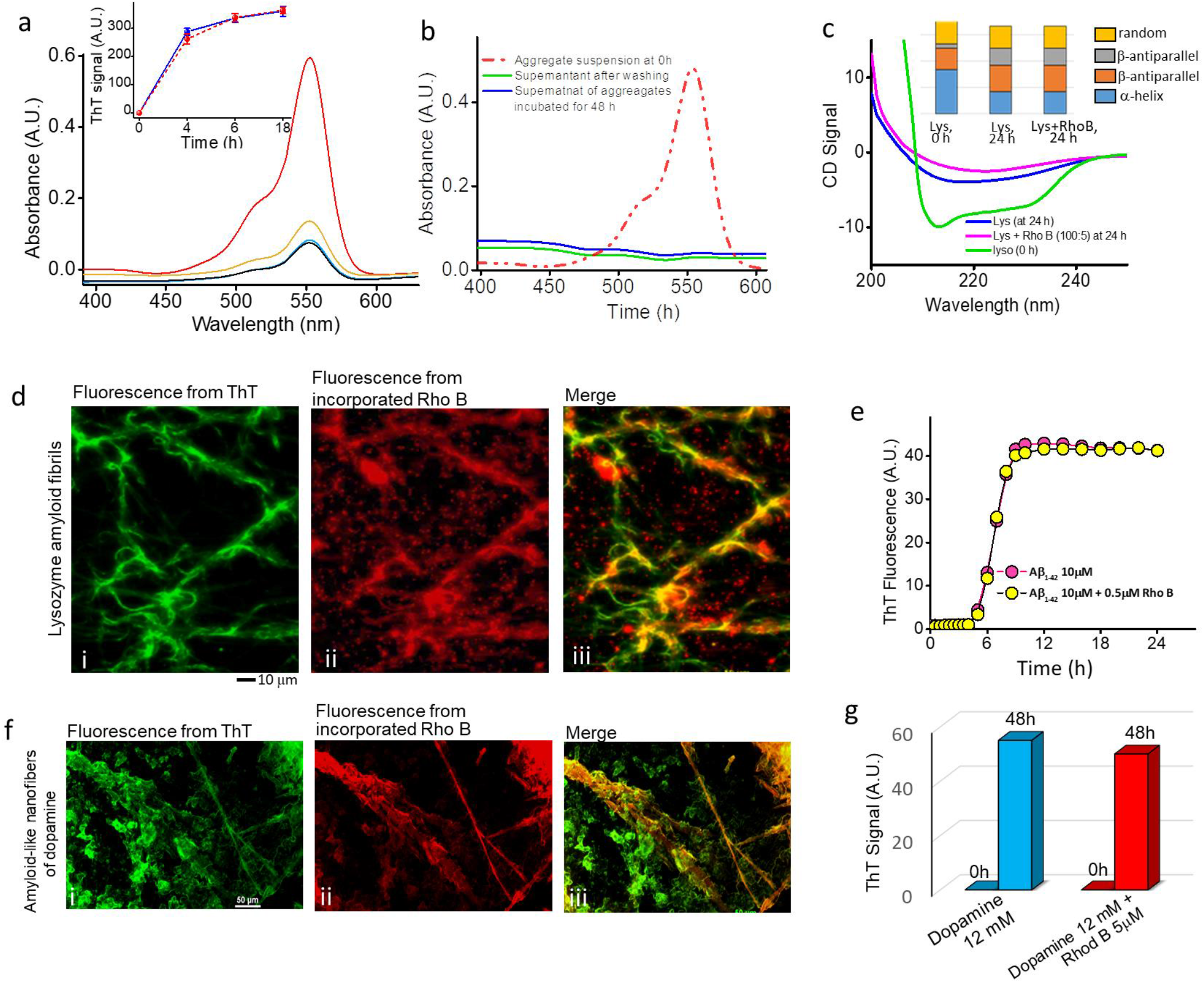
Noncovalent incorporation of Rhodamine B into Lysozyme, Aβ and dopamine fibrils. ***a,*** Decrease in the Rho-B absorbance in the supernatant of centrifuged [140 μM Lys + 7 μM Rho-B] sample during aggregation at different time points. Inset shows the ThT assays on amyloid formation of lysozyme in PBS, pH 7.4: 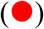 Lys alone; 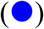 lys + Rho-B (at 100:5 molar ratio); ***b,*** Absorbance of Rho-B in (Iys+RhoB) sample at different conditions: 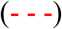 supernatant at 0 h; 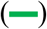 supernatant of the centrifuged resuspended [Lys+Rho-B] aggregates; 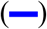 supernatant of the centrifuged [Lys+Rho-B] aggregate suspension after 2 d storage. ***c,*** CD data confirming the retained β-sheet contents of Lys amyloid aggregates after Rho-B incorporation: 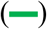 Lys at 0 h; 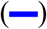 Lys aggregates at 48 h; 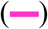 [lys +Rho-B] aggregates at 48 h. Inset shows the % predicted secondary structures, as labeled. ***d,*** Fluorescence microscopy images of the Rho-B incorporated lysozyme fibrils: (i) ThT-stained (external dye); (ii) Fluorescence of incorporated Rho-B from lysozyme fibrils; (iii) Merged view revealing the internalization of Rho-B into amyloid fibrils. ***e,*** ThT assays on amyloid formation of 10 μM Aβ_1-42_ in the presence and absence of Rhodamine B in PBS, pH 7.4: 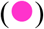 Aβ_1-42_ alone; 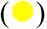Aβ_1-42_ + RhoB at 100: 5 molar ratio. ***f,*** Fluorescence microscopy images of the Rho-B incorporated dopamine fibrils: (i) ThT-stained (external dye); (ii) Fluorescence of incorporated Rho-B from dopamine fibrils; (iii) Merged view revealing the internalization of Rho-B into dopamine fibrils. ***f,*** Histograms showing ThT assays of dopamine aggregation in the presence and absence of RhoB, as labeled.

Since recent studies has shown the formation of amyloid-mimicking nanostructures via aggregation of single metabolites[16, 17, 30] we wanted to validate non-covalent incorporation RhoB into metabolite aggregates. For this we successfully made RhoB incorporated nanostructures of dopamine and our fluorescence microscopy data confirms the successful incorporation of RhoB into amyloid-like dopamine nanostructures without affecting the aggregation kinetics **(Figure 3 f-g).**

### Application of RhoB-incorporated amyloids for experiments in cell systems

Next, we extended this investigation to test whether this method can be applied for studying cellular activities of amyloid aggregates. We chose human neuroblastoma cells (SH-SY5Y cell line) because it is considered as a convenient cell model system for amyloid based assays [31]. The control SH-SY5Y cells without insulin amyloids remained unaffected (**Figure 4a**); however, severe cell damaging effect was observed when the cells were treated (at 24 h) with RhoB-incorporated insulin amyloid fibrils (**Figure 4b**). Fluorescence microscopy images clearly showed the presence of red puncta of insulin aggregates in the damaged cells (**Figure 4b, panel iv**). Cellular internalization of RhoB-incorporated insulin aggregates was further confirmed by 3D construct’s view built from z-stack confocal images as shown in **Figure 4c.** We further confirmed the internalization of the insulin aggregates by performing live cell imaging for 1.5 h after the aggregate treatment. Images shown in **Figure 4d** represents magnified snapshots captured from the video file of the live cell imaging (supplementary information). To further generalise the application of the fluorescence microscopic study of this method in cellular systems, we extended this experiment to see internalization of RhoB incorporated lysozyme aggregates into SH-SY5Y cells (Figure **5a**, **5b**). Furthermore we also captured internalization of RhoB-incorporated lysozyme aggregates into a lung cancer cell lines (A549) treated with RhoB-incorporated lysozyme aggregates (10 μM, 24 h). Data shown in **Figure 5c** and **5d** revealed similar results, as shown in **Figure 4**, supporting the wider applicability of this method in amyloid related fluorescence microscopic studies involving different types of proteins and cells.

**Figure 4.**
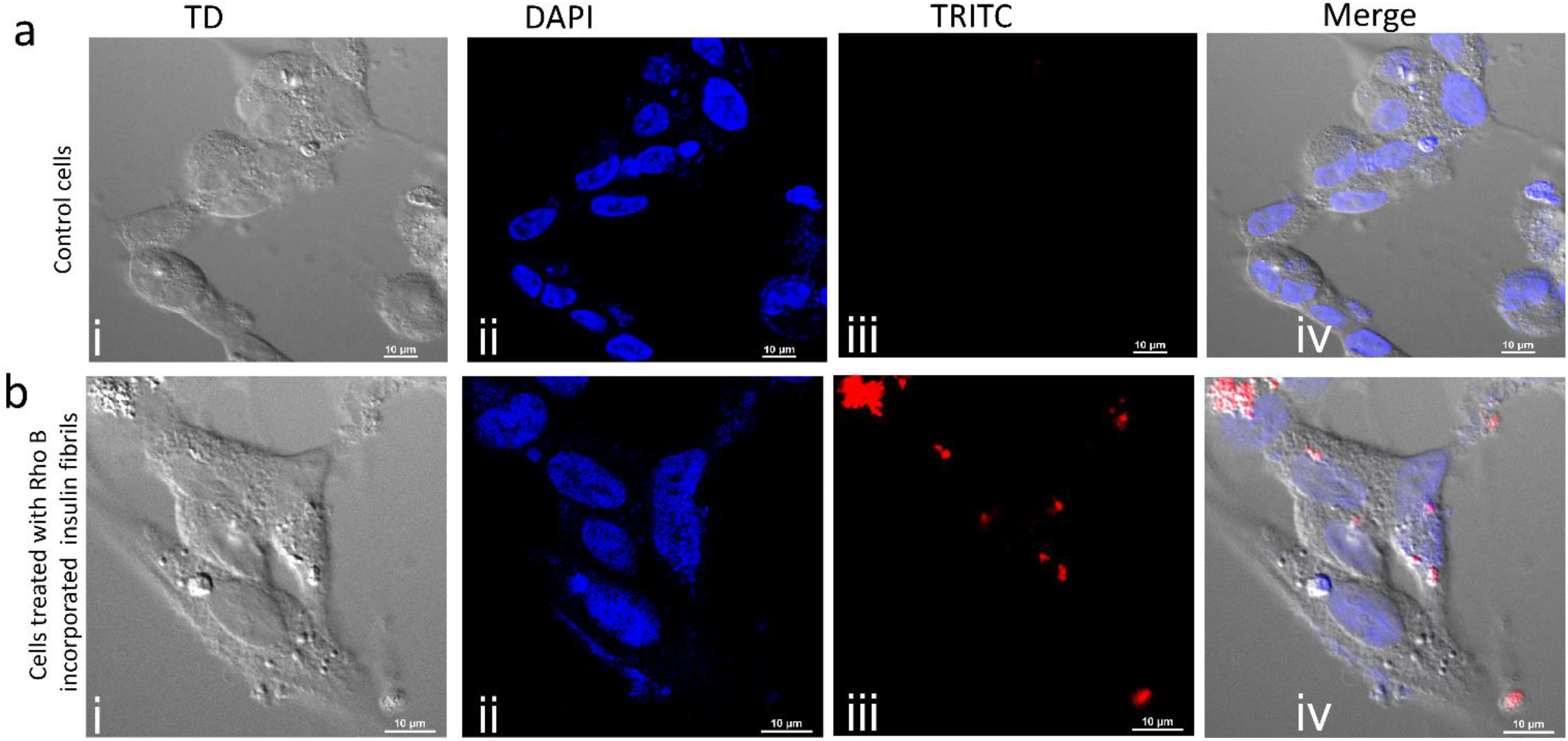
Confocal imaging of internalization of RhoB incorporated insulin aggregates into human neuroblastoma cells (SH-SY5Y cells). ***a,*** Control untreated cells visualized through: (i) TD image; (ii) nucleus staining via DAPI (blue); (iii) visualization via TRITC, indicating absence of red puncta; (iv) Merged view of (i), (ii) and (iii). ***b,*** SH-SY5Y cells treated with RhoB incorporated insulin aggregates at 24 h: (i)TD image; (ii) nucleus staining via DAPI (blue); (iii) visualization red puncta, indicating the internalized RhoB incorporated insulin aggregates; (ii) Merged view of (i), (ii) and (iii).

**Figure 5.**
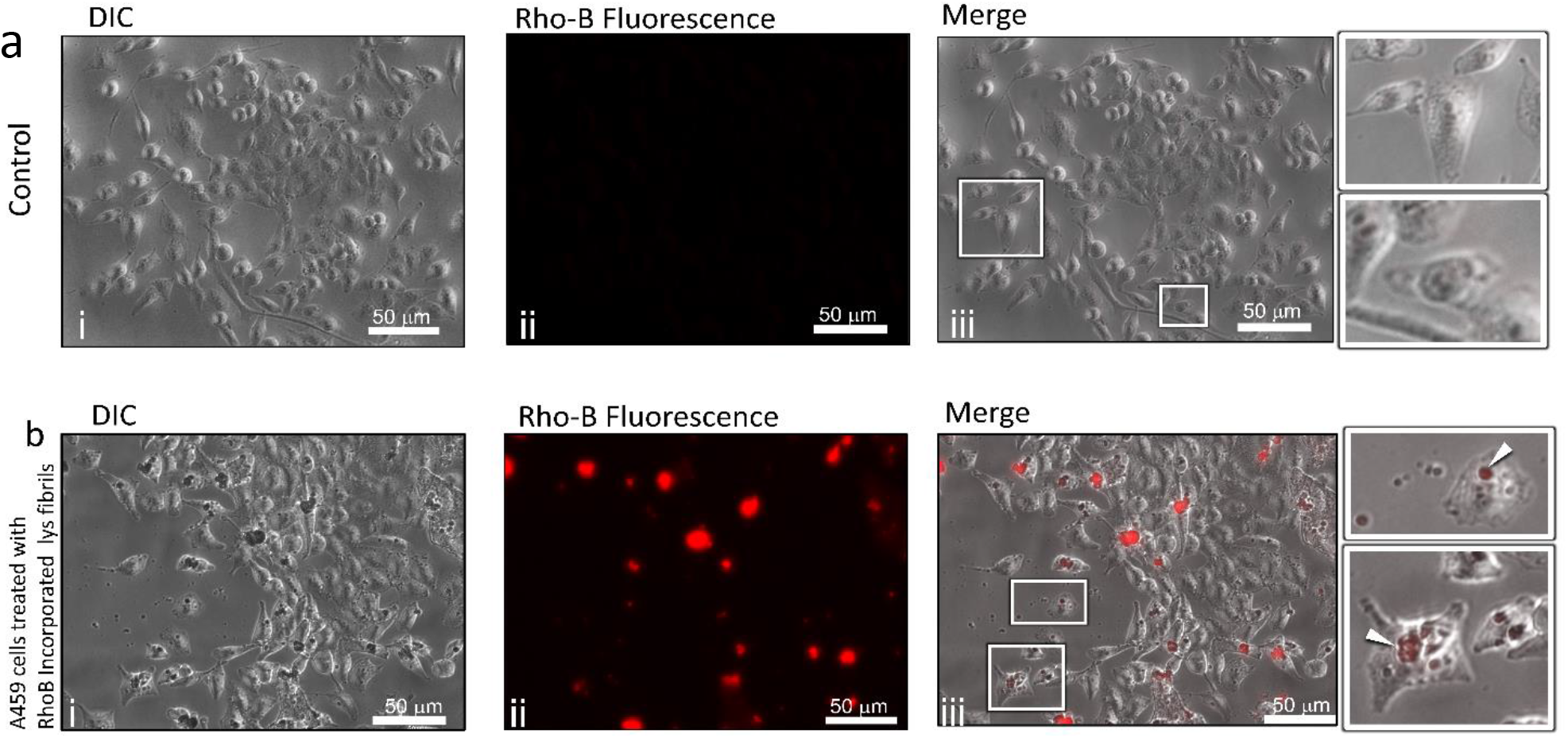
Fluorescence microscopy shows internalization of Rho-B incorporated lysozyme fibrils into cells. ***a,*** Fluorescence microscopy of untreated control A549 cells: (i) DIC image for showing intact cells. (ii) RhoB-fluorescence visualization of cells via TRITC filter; (iii) Merge of (i) and (ii). Inset showed magnified views as indicated. Scale bar for all images 50 μm. ***b,*** A549 cells treated with RhoB-incorporated lysozyme amyloid aggregates: (i) DIC image revealed presence of cells with altered morphology. (ii) RhoB-fluorescence visualization of insulin-amyloid treated cells via TRITC filter showed presence of red puncta; (iii) Merge of (i) and (ii) showed presence of red puncta in cells, confirming the internalization of RhoB-incorporated amyloid aggregates of lysozyme in cells. Insets are the magnified view of (iii). Scale bar for all images 50 μm.

It was important to examine whether RhoB alone at the studied concentrations has any cytotoxic effect, hence we performed MTT assay. The results did not show any alterations in the cell viability after RhoB alone was added to the SH-SY5Y cells, confirming the nontoxic nature of RhoB at studied concentrations (0.5-10 μM) (**Figure S4a**). Previous studied have found that RhoB is nontoxic up to 30 μM[32]. Further, our confocal data on the SH-SY5Y cells treated with only RhoB (10 μM) revealed absence of red puncta in the TRITC imaging (**Figure S4b**). Hence, RhoB alone is impermeable to SH-SY5Y cells at the studied concentrations[33, 34]. All these data suggest that fluorescence microscopy of the amyloid-induced cellular activities can be studied by preparing fluorescent amyloid fibrils via the non-covalent incorporation of RhoB, as described in this study.

### FRET studies using fluorophore incorporated amyloid fibrils

Since we successfully established the non-covalent incorporation of fluorophores into protein amyloid fibrils, we extended this study to see whether FRET experiments (methods in supplementary information) can be conducted using this method. Using the protocol, as described above, we prepared two samples of insulin fibrils: (1) RhoB incorporated insulin fibrils, (2) Fluorescein incorporated insulin fibrils. The fibrils samples were washed three times to get rid of any soluble fluorophores in the sample. Data shown in **Figure 6d-e** shows the RhoB and Flu specific emission profiles for the respective insulin fibrils. When a mixture of both insulin fibrils (Ins-Fibrils^RhoB^ + Ins-Fibrils^Flu^) was excitation at 490 nm (Fluorescein specific), two distinct emissions (RhoB and Fluorescein) were observed. Since Fluorescein and Rhodamine are known FRET pairs [25], this data suggests direct interactions between fibrillar structures of insulin. Hence, this method can be used for successful FRET detection between interfibrillar interactions in amyloid systems.

**Figure 6.**
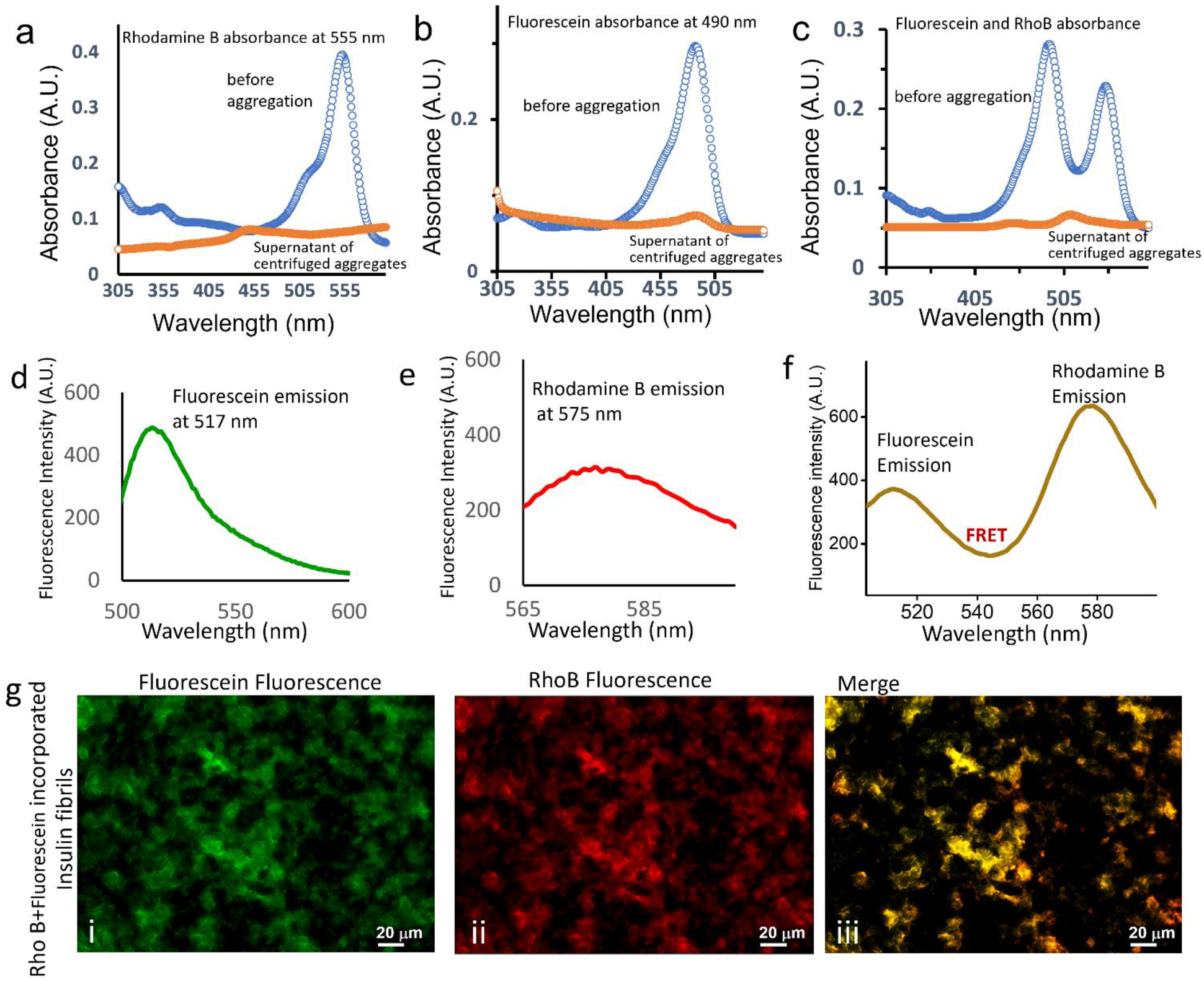
***a,*** Absorbance spectra showing the incorporation of Rhodamine B into insulin fibrils: supernatant of centrifuged [RhoB + Ins] before aggregation 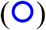 and after aggregation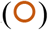. ***b,*** Absorbance spectra showing the incorporation of Fluorescein into insulin fibrils: supernatant of centrifuged [RhoB + Ins] before aggregation 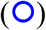 and after aggregation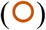. ***c,*** Absorbance spectra showing the incorporation of both Rho B and fluorescein into insulin fibrils: supernatant of centrifuged [RhoB + Fluorescein + Ins] before aggregation 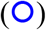 and after aggregation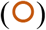. ***d,*** Fluorescence emission of incorporated Fluorescein from insulin fibrils (λ_em_ = 517nm). ***e,*** Fluorescence emission of incorporated Rho B from insulin fibrils(λ_em_ = 575nm). ***f,*** FRET signal confirms incorporation of both Rho B and Fluorescein into insulin fibrils (λ_em_ at 575 nm and 517 nm, by exciting Fluorescein). ***g,*** Fluorescence microscopy images of insulin fibrils incorporated with both RhoB and Fluorescein: (i) Fluorescein fluorescence; (ii) RhoB fluorescence; (iii) Merged view confirming the internalization of both Fluorescein Rho-B into same insulin fibrils.

Since our method established the non-covalent incorporated of fluorophores into protein aggregates, we were curious to know whether two different fluorophores can be incorporated into the same protein fibrils using our method. Hence, we examined the incorporation of both RhoB and Fluorescein into the same insulin aggregate. Both our UV-vis and fluorescence spectroscopy data confirm the simultaneous incorporation of both RhoB and Fluorescein into insulin fibrils (**Figure 6 a-e**). Next, using fluorescence spectroscopy, FRET signal was obtained by exciting the aggregates sample at

Our experimental data confirmed the successful incorporation of RhoB and Fluorescein into amyloid structures. We employed molecular docking experiments to understand the interactions between proteins and the fluorescent dye RhodB (methods in supplementary information). Both insulin and lysozyme showed strong affinity for RhoB, as evident from the analysis of docking data (**Figure 7a,7c, S1-S3**). Since RhoB is readily incorporated into amyloid structures, we extended our molecular docking study to decipher the interaction between cross-β structure of Aβ_1–42_ and RhoB. Our docking data indicated strong affinity between cross-β structure and RhoB (**Figure 7b**). Further molecular dynamics simulation for RhoB-Aβ-cross-β complex was carried out for 50 ns. RMSD (**Figure 7d**) and free energy landscape data **(Figure 7f, S5**) suggest stable complex formation between cross-β structure of Aβ1–42 RhoB. The binding free energy profile of the complex was extracted from the simulation data, as shown in **Figure 7e**.

**Figure 7.**
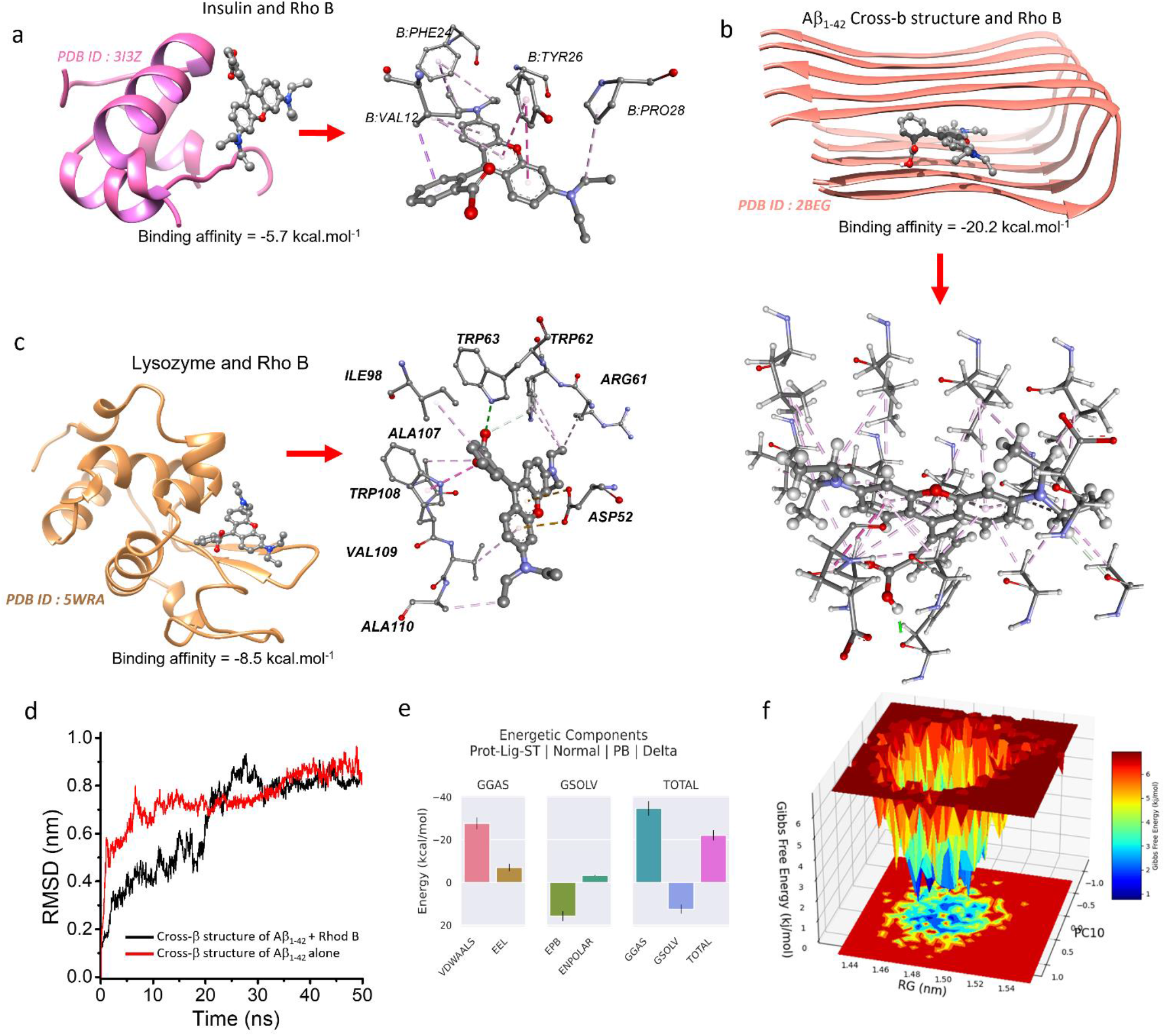
Molecular docking of RhoB with protein monomers and cross-β structures. ***a***, Docking studies on the interaction of RhoB with native structure of insulin (PDB ID: 3I3Z). Binding affinity energy = 5.7 kcal.mol^-1^. ***b***, Docking studies on the interaction of RhoB with cross-β structure of Aβ_1-42_ peptide (PDB ID: 2BEG). Binding affinity energy = −20.2 kcal.mol^-1^. ***c,*** Docking studies on the interaction of RhoB with lysozyme (PDB ID: 5WRA). Binding affinity energy = −8.5 kcal.mol^-1^. ***d,*** The RMSD value of protein ligand complex (Figure 7b) for 50 ns time stretch, as labelled. ***e***, Binding free energy profile of RhoB-Aβ_42_-fibril complex extracted from simulation data from Figure 7d. ***f,*** Free energy landscape from simulation of Rhodamine B and Aβ1-42-cross-β complex. Data for only Aβ1-42 is given in FigS5a.

## 4. CONCLUSIONS

In this work, we demonstrated an affordable and simple protocol for preparation of fluorescent amyloid aggregates of proteins. The resultant fluorophore-incorporated amyloid fibrils retained cross-β architecture which also showed amyloid-like structural and cytotoxic properties. This method also allows to conduct FRET studies using suitable donor and acceptor fluorophore pairs. Fibril disassembly experiments can also be performed by using this method via monitoring the released fluorescent dye into the solvent from the disaggregating amyloid structures (**Figure 2**). One important advantage of this protocol is that the incorporated fluorophores do not leach out during prolonged storage and after serial dilutions, and this property would greatly benefit studies involved with cellular activities of protein amyloids. Although covalent tagging of fluorophores with proteins is important for current amyloid research, this protocol is usually very expensive and full of difficulties in both the synthesis and the yield of the desired peptides. Importantly, the attached fluorescent dye, as an inbuilt moiety in the polypeptide chain, may influence the inherent properties of the proteins including their native conformation, their aggregation kinetics, and the architecture of their amyloid assembly. In contrast to the unavoidable complications linked to the covalent tagging of fluorophores, our method produces stable fluorescent amyloids without altering the aggregation kinetics and the characteristic structure of the amyloids. The findings of this work establish a simple and affordable protocol which could drastically assist amyloid researchers working on both *in vitro* and animal model systems.

## ASSOCIATED CONTENT

Supplementary information; Materials and Methods; Figures S1 to S5.

## AUTHOR INFORMATION

Corresponding Author

Correspondence and requests for materials should be addressed to K.K.

## Notes

The authors declare no competing financial interests.

## Author Contributions

The manuscript was written through the contributions of all authors. All authors have approved the final version of the manuscript. ^†^K.P.P and M.A. and have an equal contribution.

## ACKNOWLEDGMENTS

The authors thank Jawaharlal Nehru University for providing the required facilities from Advanced Instrumentation Research Facility (AIRF-JNU) and Central Instrumentation Facility-School of Life Sciences (CIF-SLS).

## SUPPLEMENTARY INFORMATION

**Figure S1.**
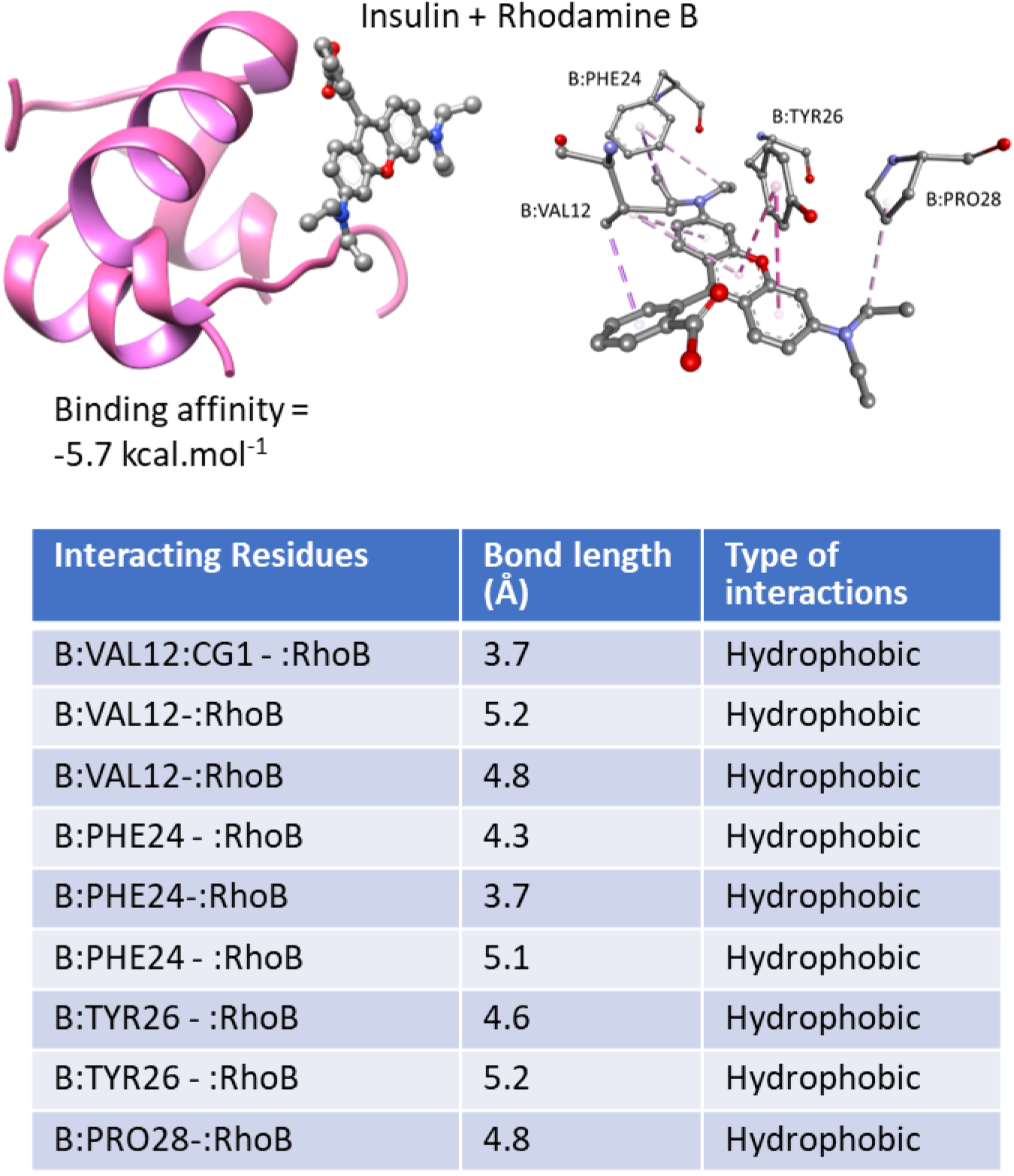
Molecular docking studies on Insulin-RhoB interaction. Table below lists the interacting residues.

**Figure S2.**
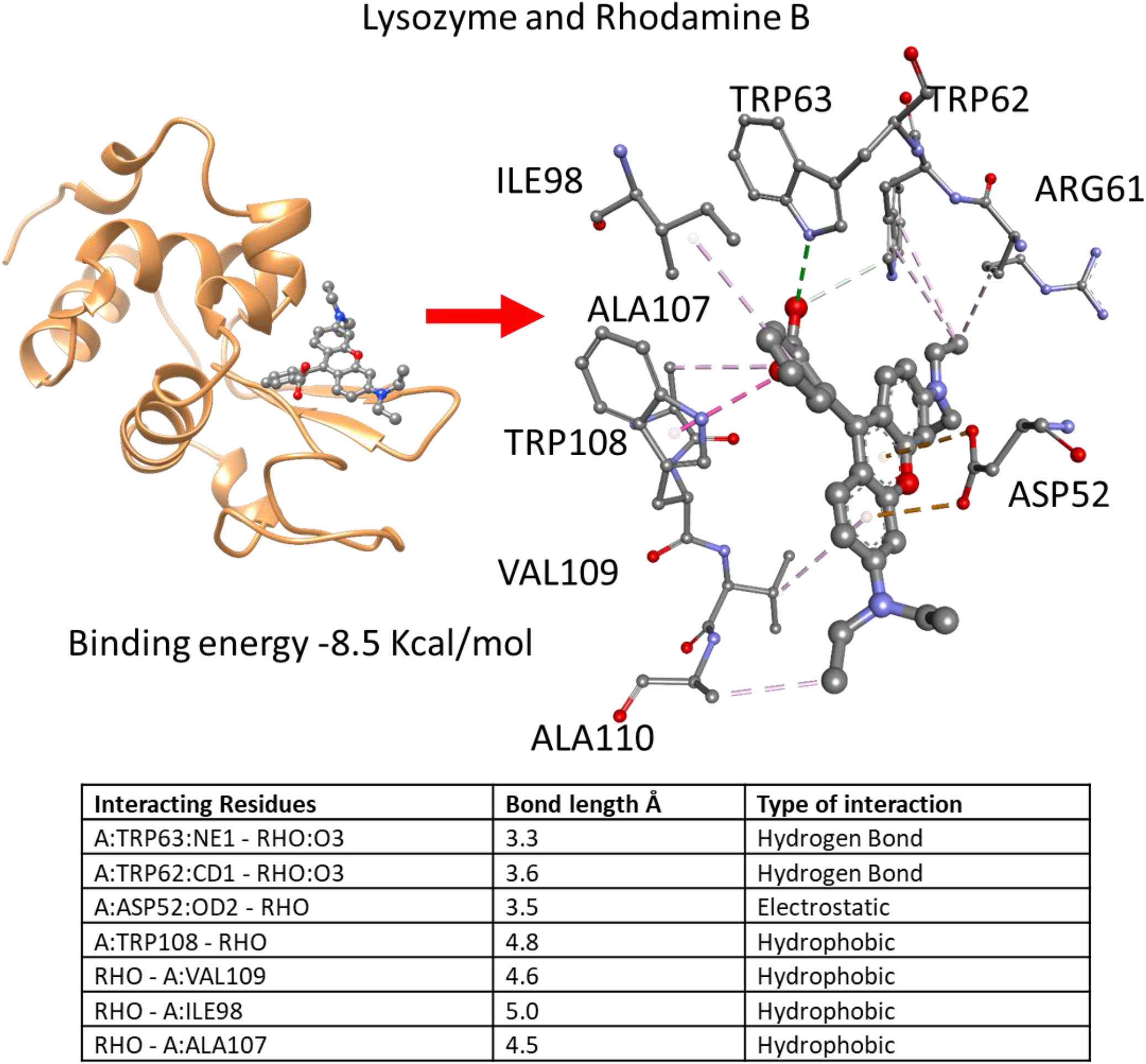
Molecular docking studies on lysozyme-RhoB interaction. Table below lists the interacting residues.

**Figure S3.**
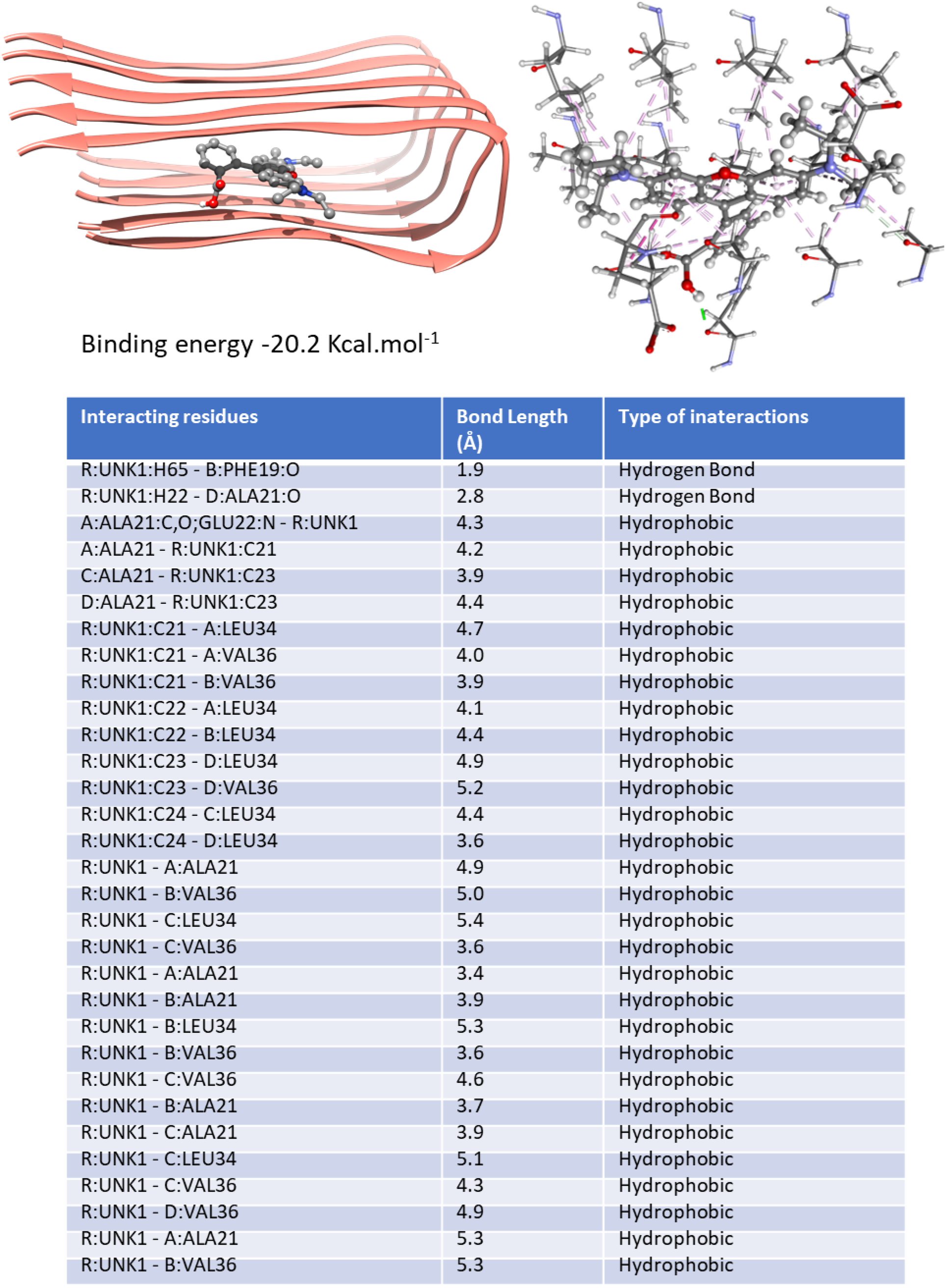
Molecular docking of cross-beta structures of Abeta1-42 peptide (PDB ID) with Rhodamine B. Binding energy −20.2 Kcal.mol^1^

**Figure S4.**
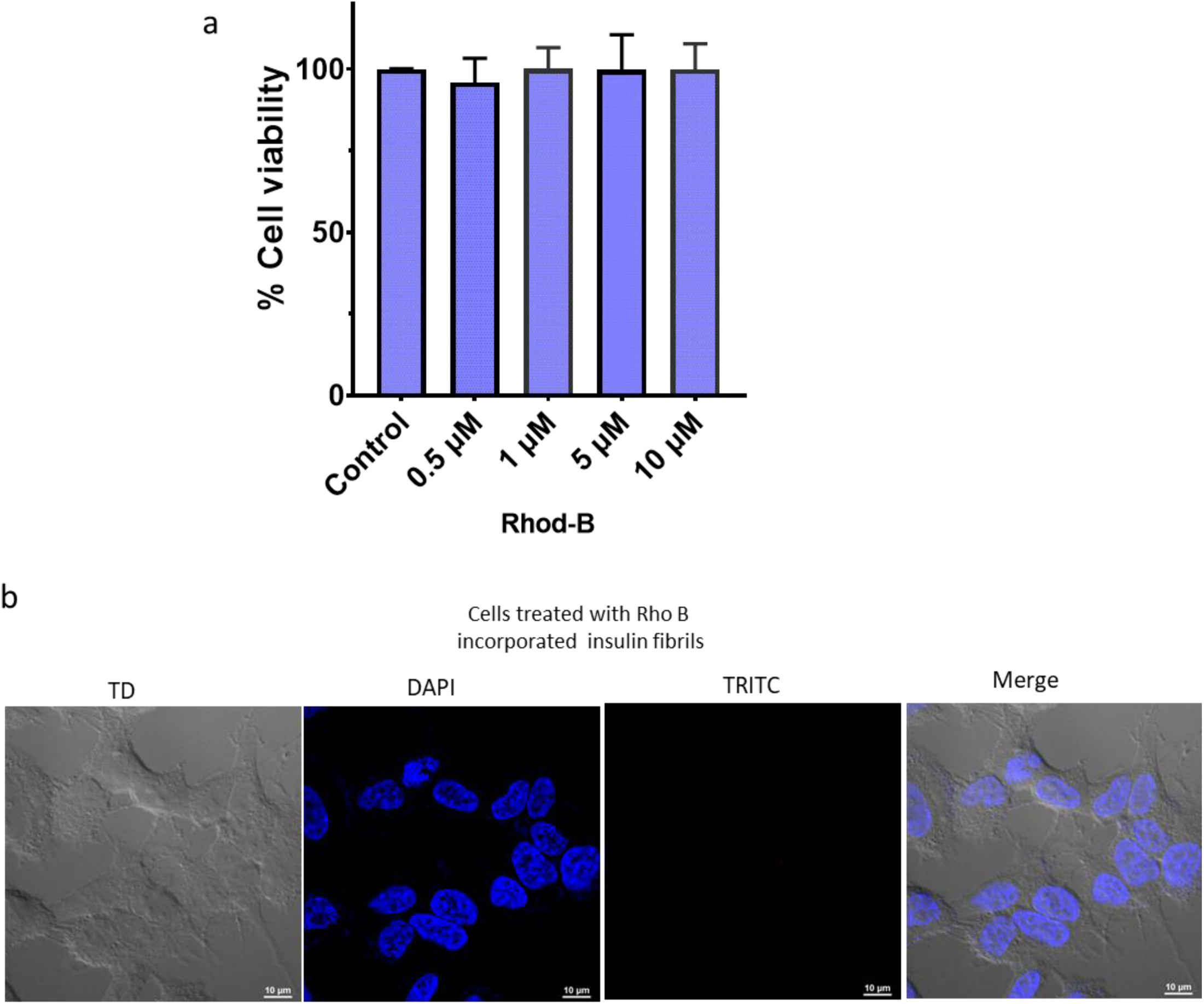
**a,** MTT assay on the effect of only Rhodamine B on SH-SY5Y cells, ***b,*** SH-SY5Y cells treated with only RhoB at 24 h: (i)TD image; (ii) nucleus staining via DAPI (blue); (iii) TRITC visualization indicating no red puncta; (ii) Merged view of (i), (ii) and (iii).

**Figure S5.**
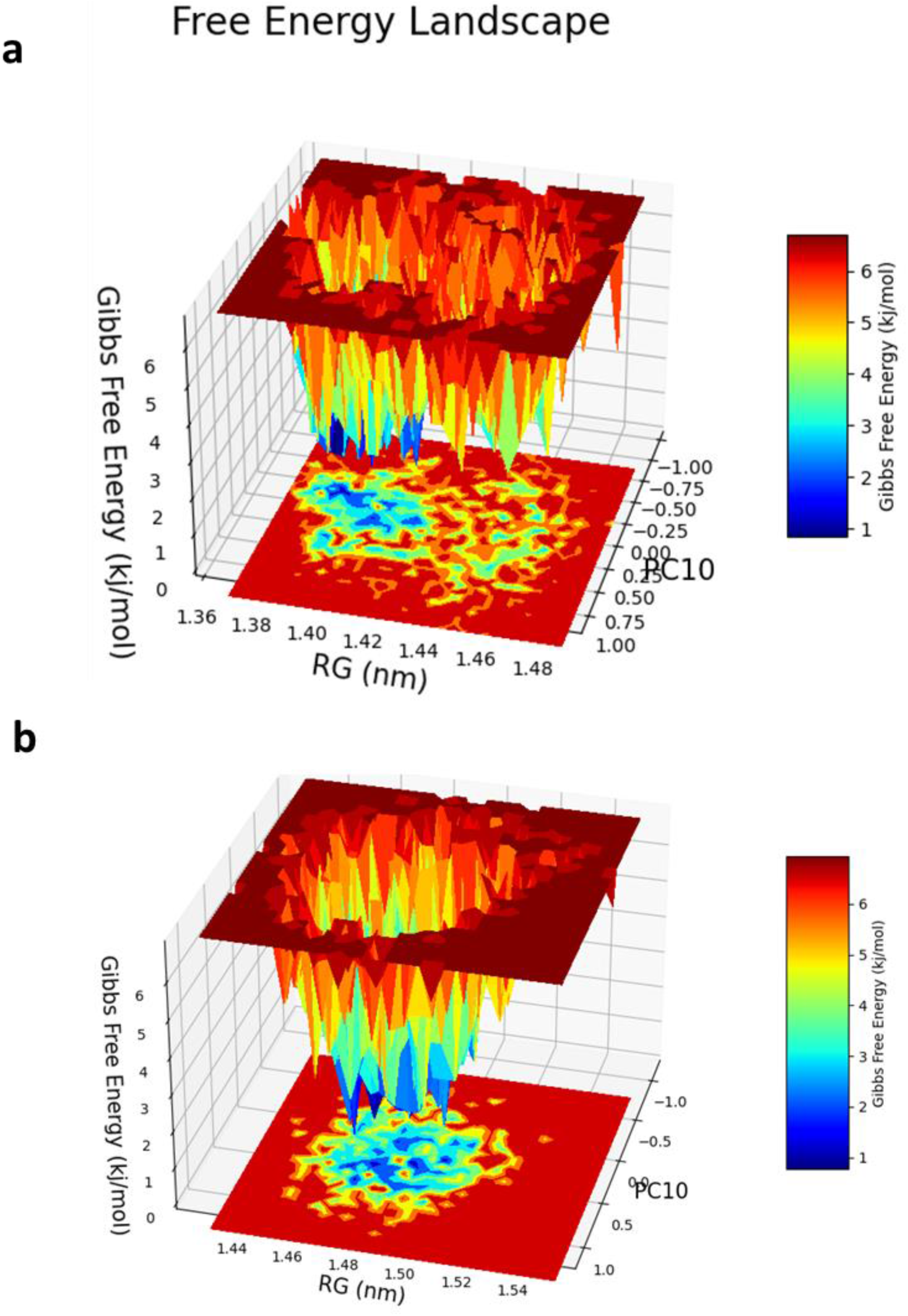
Free energy landscape from simulation of Rhodamine B and Abeta42-cross-beta complex. A, only abeta 42; b. Abeta42-Rhodamine B complex.

## METHODS

### MTT assay

The established MTT assay (*1, 2*) protocol was performed. SH-SY5Y cells were seeded in 96-well plates (5 x10 cells/well) in triplicates in the culture medium for 24 h. The confluent cells were then treated with different doses (0.5 μM, 1 μM, 5 μM and 10 μM) of Rhodamine B only samples for 24 h. Next, these cells were incubated with MTT for 4 h, and change in the color was measured using spectrophotometer at 570 nm on a microtiter plate reader (ThermoFisher VARIOSKAN microplate reader).

### Fluorescence microscopy and live cell imaging

Internalization of non-covalently RhoB tagged amyloid fibril was monitored by fluorescence microscopy. Cells (SH-SY5Y and A549) were grown in six-well plates containing glass coverslips. When cells become confluent about 70% then the cells were treated with 5 and 10 μM of RhoB tagged amyloid fibrils. Treated and non-treated cells were incubated in CO_2_ incubator at 37 °C for 24h. After 24 h the cells were washed with 2 mL of PBS followed by fixation of the cells with 2 mL of 4 % paraformaldehyde for 30 min at RT. Next, paraformaldehyde was removed, and cells were washed three times with 2 mL of PBS. Further, the SH-SY5Y cells were mounted with Vectasheild with DAPI (Vector Laboratories H-1200) on glass slides. Nikon AIR HD Fluorescent Confocal Microscopy was used to visualize the internalized RhoB-tagged amyloid aggregates and their uptake by the cells, Z-stack images of internalized fluorescent amyloid fibrils in cells were captured at 60 x. To further monitor the internalization process of these fluorescent labelled amyloid fibril we performed the live cell imaging of SH-SY5Y cells treated with RhoB-incorporated insulin aggregates by using confocal microscopy, cells were grown on confocal dish with glass bottom in CO_2_ incubator, till the 70-80% confluency. Next, we monitored the internalization for 1.5 h after the treatment in live cell via confocal microscopy and each snapshot was captured in every 10 s. Inbuilt NIS element software was used for data acquisition, image analysis and live cell imaging. We also visualized A549 cells treated with RhoB-incorporated lysozyme aggregates. After fixation and washing step, the cells were mounted on glass slide. The images were captured at 20 x under Nikon TiE fluorescence microscope, analysed by in-built NIS4.00.00 software.

### Circular Dichroism (CD)

The structural changes during the conversion of soluble protein monomers into beta-sheet rich amyloid aggregates were studied using Chirascan™ qCD, attached to a Peltier temperature controller. PBS was used as the reference solution and the CD spectra were recorded at room temperature using a cuvette of 2 mm path length. Data presented in the text are the average of three independent measurements.

### Fluorescence resonance energy transfer (FRET) assay

We performed FRET experiment using Shimazdu fluorescence spectrophotometer (RF-5300, Japan. For this specific assay two known FRET pair fluorophore dyes which are rhodamine B (acceptor) and fluorescein (donor) were selected. Both the dyes were incubated with soluble insulin at 65 °C for 72 hours maintaining the concentration at 1:20 molar ratio of dye:protein. After incubation the sample was subjected to centrifugation (15000 rpm) and washing steps three times to prepare the suspension of fluorophore incorporated protein aggregates. Next, using fluorescence spectroscopy, FRET signal was obtained by exciting the sample at donor’s excitation wavelength (490nm). Two distinct emission peaks (specific for RhoB and fluorescein) were recorded.

### Molecular docking studies

Molecular docking study was conducted by Autodock Vina using PyRx open-source software (GUI version 0.8)(*3, 4*). We obtained the structure of Rhodamine B from PubChem (https://pubchem.ncbi.nlm.nih.gov/) (PubChem <u>CID:6694</u>). PDB structures of the studied proteins were obtained from RCSB (Lysozyme PDB ID: 5WRA(*5*), Insulin PDB ID 3I3Z(*6*), Aβ_42_ fibril PDB ID: 2BEG (*7*)). Prior to docking, pre-processing and protonation of receptor molecules were performed by removing the unwanted ligand and water molecules and by adding polar hydrogen atoms. For docking, AutoDock Vina used a search space of grid box size (x = 24 Å, y = 24 Å, z = 24 Å) for 3I3Z, (x = 24 Å, y = 24 Å, z = 24 Å) for 5WRA and (x = 22 Å, y = 13 Å, z = 16 Å) for 2BEG. The results obtained were analyzed based on the binding affinity energy (kcal mol^-1^) parameters linked to the respective protein–ligand complexes. The protein–ligand docked complex with the lowest energy was chosen for further analysis. Docked complexes were visualized and analyzed by using Discovery Studio visualizer (v 16.1.0153) and Chimera 1.15.

### Molecular dynamics simulations

We performed MD simulations using the GROMACS 2020.6 (https://doi.org/10.5281/zenodo.4576060)(*8*) after docking of protein and RhoB. Here, we performed MD simulation of the complex between amyloid beta fibril and RhoB. Topologies for protein and protein–ligand complexes were produced using the CHARMM 36 force field(*9*). The complex and single protein structures were solvated in the water model after the topology file was created, and structures were neutralized by adding ions. After that, these structures were relaxed using an energy minimization approach involving the steepest descent Algorithm and the Verlet cut-off scheme that was run for 50,000 cycles at 10 kJ/mol. The equilibration step of protein and ligands complex was performed on NVT (constant volume) as well as NPT (constant pressure) for 1000 ps trajectory period. After equilibration step, the simulation analysis was calculated at 300 K temperature and 1 atm pressure using 2 fs time step for a 50 ns. The obtained trajectory files were used to visualize the deviation of protein and complex to determine the system’s stability in a water environment. To investigate the deviation between protein and ligand complexes we use Root mean square variance (RMSD), Radius of gyration (RG), and Principal component analysis (PCA). Further, we calculated the interaction energy between protein and ligands to calculate the strength between protein and ligand. Furthermore, Molecular Mechanics Poisson–Boltzmann Surface Area (gmx_MMPBSA)(*10*) method was used to calculate the total binding free energy (equation 1) using gmx_mmpbsa package (*11*) in GROMACS software, the free solvation energy (polar + non-polar solvation energies), and potential energy (electrostatic + Van der Waals interactions) of each protein–ligand complex for last 100 frames

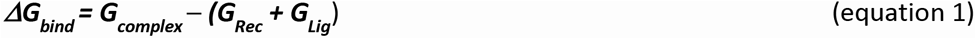

The free energy landscape of protein folding on the RhoB bound complex was measured using geo_measures v 0.8. Geo_measures include a powerful library of g_sham and form the MD trajectory against RMSD and Radius of gyration (Rg) energy profile of folding recorded in a 3D plot using matplotlib python package (*12*).

